# Manipulation of IRE1-dependent MAPK signaling by a Vibrio agonist-antagonist effector pair

**DOI:** 10.1101/2020.09.01.278937

**Authors:** Nicole J. De Nisco, Amanda K. Casey, Mohammed Kanchwala, Alexander E. Lafrance, Fatma S. Coskun, Lisa N. Kinch, Nick V. Grishin, Chao Xing, Kim Orth

## Abstract

Diverse bacterial pathogens employ effector delivery systems to disrupt vital cellular processes in the host (*1*). The type III secretion system 1 of the marine pathogen, *Vibrio parahaemolyticus*, utilizes the sequential action of four effectors to induce a rapid, pro-inflammatory cell death uniquely characterized by a pro-survival host transcriptional response (*2, 3*). Herein, we show that this pro-survival response is caused by the action of the channel-forming effector VopQ that targets the host V-ATPase resulting in lysosomal deacidification and inhibition of lysosome-autophagosome fusion. Recent structural studies have shown how VopQ interacts with the V-ATPase and, while in the ER, a V-ATPase assembly intermediate can interact with VopQ causing a disruption in membrane integrity. Additionally, we observe that VopQ-mediated disruption of the V-ATPase activates the IRE1 branch of the unfolded protein response (UPR) resulting in an IRE1-dependent activation of ERK1/2 MAPK signaling. We also find that this early VopQ-dependent induction of ERK1/2 phosphorylation is terminated by the VopS-mediated inhibitory AMPylation of Rho GTPase signaling. Since VopS dampens VopQ-induced IRE1-dependent ERK1/2 activation, we propose that IRE1 activates ERK1/2 phosphorylation at or above the level of Rho GTPases. This study illustrates how temporally induced effectors can work as in tandem as agonist/antagonist to manipulate host signaling and reveal new connections between V-ATPase function, UPR and MAPK signaling.

**Importance:** *Vibrio parahaemolyticus (V. para)* is a seafood-borne pathogen that encodes two Type 3 Secretion Systems (T3SS). The first system T3SS1 is thought to be maintained in all strains of *V. para* to to maintain survival in the environment, whereas the second sytem T3SS2 is linked to clinical isolates and disease in humans. Herein, we find that first system targets evolutionarily conserved signaling systems to manipulate host cells, eventually causing a rapid, orchestrated cells death within three hours. We have found that the T3SS1 injects virulence factors that temporally manipulate host signaling. Within the first hour of infection, the effector VopQ acts first by activating host surval signals while diminishing the host cell apoptotic machinery. Less than an hour later, another effector VopS reverses activation and inhibition of these signaling systems ultimately leading to death of the host cell. This work provides example of how pathogens have evolved to manipulate the interplay between T3SS effectors to regulate host signaling pathways.

## Introduction

The seafood-borne pathogen, *Vibrio parahaemolyticus* (*V. para*), uses two needle-like type III secretion systems (T3SS1 and T3SS2) to inject effectors into host cells to manipulate signaling and cellular processes during infection (*2*). The *V. para* T3SS2 is found in clinical isolates, is linked to disease in humans and has been shown to mediate invasion into mammalian host cells (*4, 5*). By contrast, the T3SS1 is present in all *V. para* isolates and is thus believed to be essential for survival in its environmental niche. This niche has been rapidly expanding due to the warming of coastal waters contributing to the resurgence of *V. para* as a significant cause of gastroenteritis world-wide (*6, 7*). Together, the *V. para* T3SS1 effectors orchestrate a temporally regulated non-apoptotic cell death in cultured cells (*2*). The specific cell type that T3SS1 has evolved to target in the environment remains undefined; however, its effectors target processes that are conserved from yeast to humans (*2, 8-10*).

Cell death mediated by T3SS1 occurs in distinct and highly reproducible stages through the temporal action of four known effectors: VopQ (VP1680), VPA0450, VopS (VP1686) and VopR (VP1683) (Fig. 1A) (*2, 11*). Within 30 minutes of a synchronized infection, VopQ forms an outward-rectifying channel in V-ATPase-containing membranes resulting in neutralization of the compartment (e.g. vacuole or lysosome) and inhibition of membrane fusion (Fig. 1A) (*9, 12*). These two activities inhibit autophagic flux, resulting in massive autophagosome accumulation and contribute to a pro-inflammatory cell death within three hours(*13*). VPA0450 is a phosphatidyl 5-phosphatase that hydrolyses PI(4,5)P_2_ at about one hour after infection, resulting in blebbing of the plasma membrane (Fig. 1A) (*14*). Soon after, VPA0450-mediated blebbing is observed, VopS, a Fic (filamentation induced by cAMP) domain–containing protein, covalently attaches an adenosine monophosphate (AMP) to a threonine residue in the switch 1 region of Rho guanosine triphosphatases (GTPases) Rho, Rac and Cdc42. This modification, termed AMPylation, inactivates the Rho GTPases thereby precipitating cytoskeletal collapse and cell rounding, as well as inactivation of nuclear factor-kappaB (NF-κB) and mitogen activated protein kinase (MAPK) signaling pathways (Fig. 1A) (*10, 15*). The fourth effector, VopR, causes cell rounding around 90 minutes post infection, but its activity has remained elusive (*11*). Despite the rapid, non-apoptotic cell death orchestrated by the T3SS1, our previous studies have uncovered evidence that the T3SS1 rewires host gene expression to subvert cell death and activate cell survival pathways, including MAPK signaling pathways. We performed a systems level analysis of the temporal changes in host cell gene expression during *V. para* infection to understand how the T3SS1 effectors work in concert to orchestrate cell death and subvert host immune responses (*3*). The host transcriptional response to T3SS1 resulted in the activation of host cell survival networks and repression of cell death networks (*3*).

**Figure 1.**
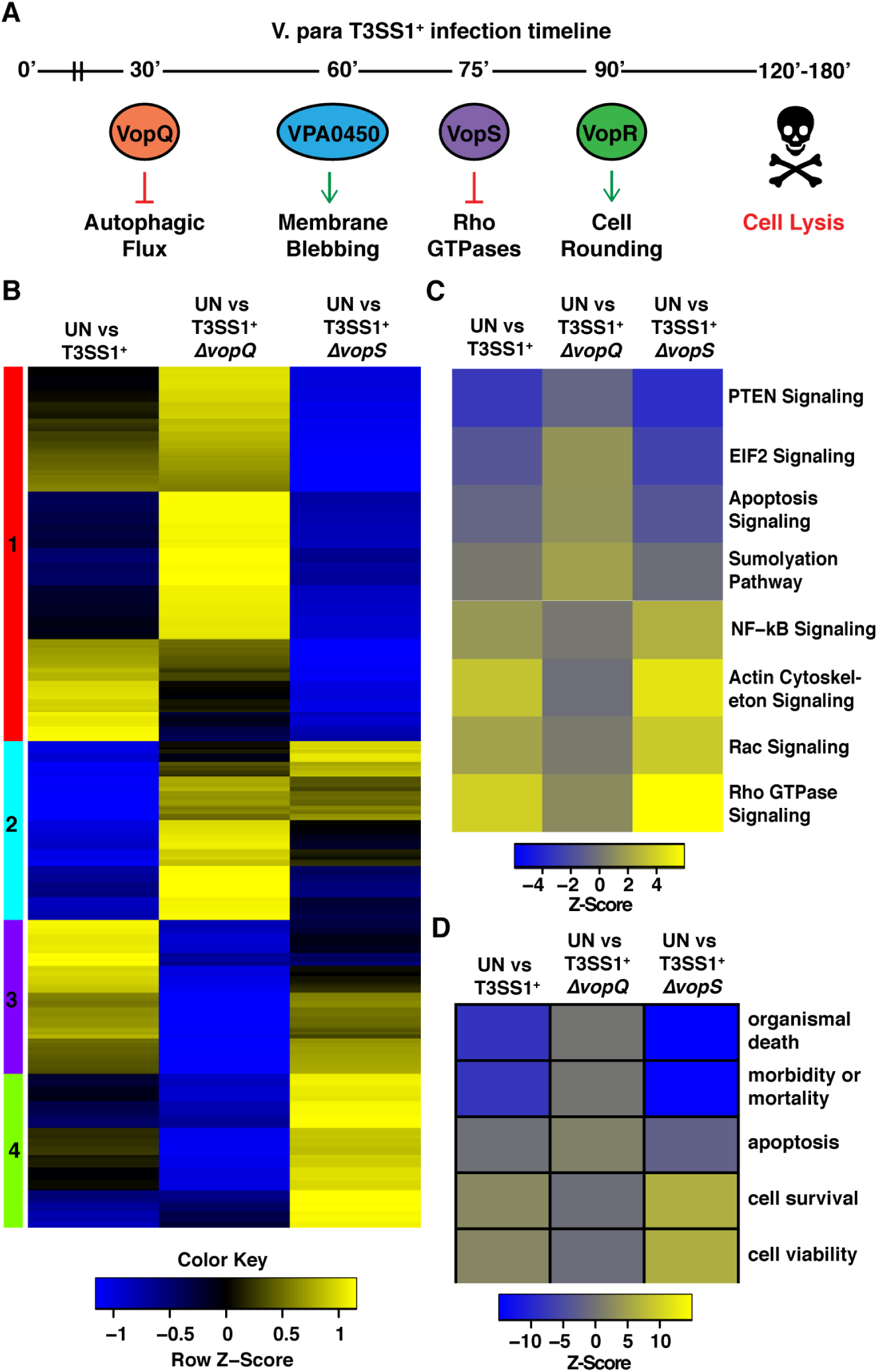
VopQ and VopS have antagonistic effect on T3SS1-specific pathway and network induction. **A)** Illustration of temporal effector function during T3SS1-mediated cell death **B)** Heatmap of normalized differential expression of previously identified T3SS1-specific transcripts in uninfected (UN) primary human fibroblasts compared to primary human fibroblasts infected with either *V. para* T3SS1^+^, T3SS1^+^ Δ*vopQ*, or T3SS1^+^ Δ*vopS* for 90 minutes. Yellow denotes transcripts with relative increased abundance infected cells compared to UN cells, and blue denotes a decreased abundance. Clusters (color bars on the left) were assigned through hierarchical clustering of the differential expression data. **C)** Heatmap of predicted repression (blue) and activation (yellow) Z-scores calculated from differential expression data for UN vs. *V. para* T3SS1^+^, UN vs. *V. para* T3SS1^+^ Δ*vopQ*, and UN vs. *V. para* T3SS1^+^ Δ*vopS* using QIAGEN’s Ingenuity^®^ Pathway Analysis software. The color key correlates the displayed heatmap color and calculated Z-scores and gray denotes unaffected (*p*>0.05) pathways. **D)** Heatmap of Ingenuity^®^ Pathway Analysis Z-score prediction of repression (blue) or activation (yellow) of biological networks after 90 minutes of POR3:T3SS1^+^, T3SS1^+^ Δ*vopQ*, and T3SS1^+^ Δ*vopS* infection.

Previously, it was found that VopQ was both necessary and sufficient for the accumulation of LC3-positive autophagosomes as well as the deacidification of endolysosomal compartments. These effects are caused by direct interaction between VopQ and the V_o_ subcomplex of the V-ATPase. Recently, cryo-EM studies performed in our lab revealed extensive interactions between VopQ and the c subunit of V_o_ V-ATPase that stabilize the insertion of VopQ in the membrane alongside the V-ATPase (*16*). The interactions of VopQ with the c-ring of V_o_ is predicted to form an unconventional membrane pore through the juxtaposition of charged resides of VopQ against the hydrophobic lipid environment (*16*). This disruption is predicted to lead to the deacidification of the lysosomal membrane. Remarkably, a fully functioning or assembled V-ATPase at the vacuole is not necessary to induce VopQ toxicity in yeast. We found that VopQ can interact with an assembly intermediate of the V-ATPase (V_o_ c-ring) in the ER resulting in cell death (*16*). As VopQ forms a pore in target membranes, the ER membrane is compromised, and this could lead to the induction of host cell signaling events including the unfolded protein response (UPR).

Herein, we find that VopQ activates the IRE1 branch of the UPR in yeast and cultured cells. We demonstrate that the activation of IRE1 by VopQ results in an activation of extracellular singal-regulated protein kinase 1/2 (ERK1/2) signaling that is dependent on IRE1 kinase but not nuclease activity. We also find that another T3SS1 effector VopS dampens VopQ-mediated activation of ERK1/2 signaling by AMPylation-dependent inactivation of Rho GTPases, thereby limiting the activation of ERK1/2 signaling to early infection time points. Taken together, our work provides another example of the powerful interplay between T3SS effectors and how they can temporally regulate host signaling pathways.

## Results

### VopQ and VopS have antagonistic effect on T3SS1-specific pathway and network induction

Previously, we discovered that the T3SS1 activates host cell survival networks and represses of cell death networks. Since, autophagy is linked to pro-survival network signaling, our findings led us to ask if VopQ could be responsible for pro-survival signals observed during infection. To understand the contribution of individual effectors to the T3SS1-specific transcriptional response, we characterized infection of primary human dermal fibroblasts (PHDFs) with *V. para* strains carrying single-effector deletions for either *vopQ* or *vopS*. We chose this cell line so that our data would be comparable with the data obtained in our previous transcriptomic analysis (*3*). We included VopS because it targets Rho GTPases that regulate MAPK signaling (*15, 17*). We used *V. para* strain, POR3, a derivative of the clinical strain RIMD2210633 that does not produce functional hemolysins or a functional T3SS2 (Δ*tdhAS*Δ*vcrD2)*, but maintains an active T3SS1 (*6*). This strain and its Δ*vopQ* and Δ*vopS* derivatives will be referred to herein as T3SS1^+^, T3SS1^+^ Δ*vopQ* and T3SS1^+^ Δ*vopS*, respectively (Table S1). As observed with previously characterized cell types, cytotoxicity of PHDFs occurring within the first four hours of infection was completely dependent on VopQ (Fig. S1A,B) (*2*). We then performed RNA-sequencing on the PHDFs after 90 minutes of infection with *V. para* T3SS1^+^, T3SS1^+^ Δ*vopQ* and T3SS1^+^ Δ*vopS* strains. The sequencing data passed statistical quality control tests, and principal component analysis indicated tight clustering of replicates (Fig. S2A). Complete differential expression data is reported in Table S2, but for this study we focused on the 398 host genes previously found to be differentially expressed specifically in response to the T3SS1 (*3*).

The hierarchically clustered expression heatmap in Fig. 1B illustrates how the T3SS1 causes changes in expression of these 398 genes in the absence of either VopQ or VopS. Of these T3SS1-specific genes, 146 were similarly differentially expressed in the uninfected (UN) vs. T3SS1^+^ and UN vs. T3SS1^+^ Δ*vopQ*-infected cells, and 197 were similarly differentially expressed in the UN vs. T3SS1^+^ and UN vs. T3SS1^+^ Δ*vopS*-infected cells (Fig. S2B, Table S3). 252 and 201 T3SS1-specific genes were either not differentially expressed or changed direction during T3SS1^+^ Δ*vopQ* and T3SS1^+^ Δ*vopS* infection, respectively. Expression of many genes, especially those within clusters 1 and 4, was oppositely affected during infection with T3SS1^+^ Δ*vopQ* compared to during infection with T3SS1^+^ Δ*vopS* (Fig. 1B, Table S3). Notably, expression of the *EGR1* and *FOS* transcription factors, which are known to be regulated by MAPK signaling pathways, was reduced in T3SS1^+^ Δ*vopQ*-infected cells compared to T3SS1^+^ -infected cells, and highly elevated by T3SS1^+^ Δ*vopS* infection (Table S3)(*18*). We validated these findings by quantitative RT-PCR and using *V. para* strains deleted for multiple effectors (T3SS1^+^ Δ*vopQR*Δ*vpa0450* and T3SS1^+^ Δ*vopRS*Δ*vpa0450*) and showed that VopQ is necessary and sufficient for the elevated expression of both *FOS* and *EGR1* (Fig. S2C,D).

We next used Ingenuity Pathway Analysis (IPA) to understand how the activities of VopQ and VopS contribute to the changes in host signaling events induced by the T3SS1 (Table S4). The T3SS1-specific induction or repression of many pathways was dependent on VopQ and enhanced in the absence of VopS (Fig. 1C). For example, induction of NF-kB signaling, actin cytoskeleton signaling and Rho GTPase signaling by T3SS1 was greatly reduced in the absence of VopQ and enhanced in the absence of VopS. These observations are consistent with the opposing effects on differential expression patterns observed in Fig. 1B.

### VopQ induces pro-survival signaling networks

To understand the relative contributions of VopQ and VopS to the host response to T3SS1 on the network level, we used IPA to perform biological function network analysis. Previously, we had shown that the T3SS1 activates cell survival networks and represses cell death networks(*3*). Strikingly, this effect was completely lost during T3SS1^+^ Δ*vopQ* infection and amplified during T3SS1^+^ Δ*vopS* infection (Fig. 1D, Table S5). Specifically, we observed a loss in cell survival and viability signaling network activation and death and mortality signaling network repression in PHDFs infected with *V. para* T3SS1^+^ Δ*vopQ* compared to *V. para* T3SS1^+^ while infection with T3SS1^+^ Δ*vopS* instead amplified these signaling changes (Fig. 1D). Interestingly, the apoptosis signalling network, which is normally repressed during T3SS1^+^ infection, was activated during infection with T3SS1^+^ Δ*vopQ* (Fig. 1D). These data strongly a model in which the activity of VopQ elicits significant transcriptional changes in the host cell that result in the activation of cell survival and repression of cell death networks and that VopS functions to dampen this response.

### VopQ induces a pulse of ERK1/2 signaling that is dampened by VopS

To further dissect VopQ’s effect on host signaling pathways in mammalian cells, we continued with a more genetically tractable model, mouse embryonic fibroblasts (MEFs). We characterized the cytotoxicity of the *V. para* T3SS1^+^ strain and its derivates in MEFs. The cell death induced by the T3SS1 occurred over a similar timescale in MEFs as in PHDFs and was similarly dependent on VopQ (Fig. S3A). This result was expected because the mechanism of T3SS1-mediated cell death is conserved across diverse cell types (*2, 3, 9, 15*). We chose to examine the ERK1/2 signaling pathway because the RNA-sequencing data suggested that VopQ activates Rho GTPase signaling (Fig. 1C) and in previous work we demonstrated that T3SS1-induced *EGR1* and *FOS* expression requires active MEK1/2, the kinase upstream of ERK1/2 (*3*). Furthermore, as we did not observe agonist and antagonist effects of VopQ and VopS, respectively, on the expression of c-Jun N-terminal kinase (JNK) signaling target genes, we focused our analysis on the ERK1/2 pathway (Table S6).

To test if VopQ induces *EGR1* and *FOS* expression by activating the ERK1/2 MAPK pathway early during infection, we starved MEFs to remove basal ERK1/2 phosphorylation, infected them with *V. para* T3SS1^+^ or *V. para* T3SS1^-^ for 45, 60, 75 and 90 minutes, and probed for phospho-ERK1/2 as well as for the presence of downstream Egr1 by Western blot. We found that the T3SS1 induced a pulse of ERK1/2 phosphorylation that peaked around 45 minutes and had completely disappeared by 90 minutes post infection (Fig. 2A). Total Egr1 protein levels began to rise 60 minutes post infection and reached their maximum at 90 minutes post infection, which is an expected pattern of expression when taking into account the time for transcription and translation of the induced *Egr1* gene (Fig. 2B). Using the MEK1/2 inhibitor U0126, we found that T3SS1-induced *Egr1* expression was similarly dependent on MEK1/2 activity in MEFs as previously reported in PHDFs (Fig. S3B) (*3*).

**Figure 2.**
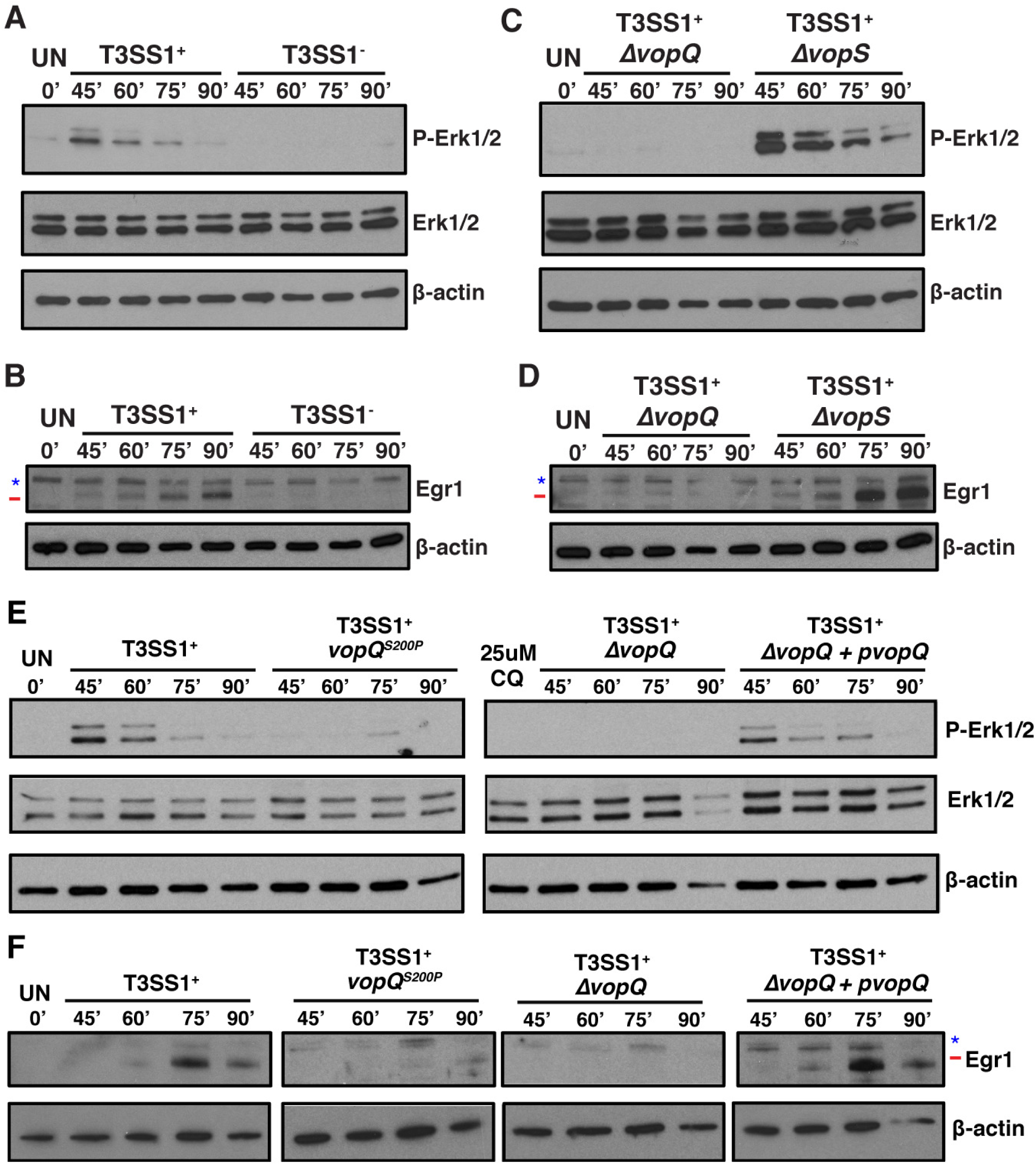
VopQ but not VopQ^S200P^ induces an early activation of ERK1/2 MAPK signaling. **A)**Immunoblot showing phosphorylated Erk1 and Erk2 (p-Erk1/2) and total Erk1/2 in starved mouse embryonic fibroblasts (MEFs) 45, 60, 75 and 90 minutes post-infection with T3SS1^+^ or T3SS1^-^ *V. para*. A pulse of p-Erk1/2 was observed early during infection with T3SS1^+^ but not T3SS1^-^ *V. para*. **B)** Immunoblot for total Egr1 in starved MEFs 45, 60, 75 and 90 minutes post-infection with POR3:T3SS1^+^ or POR3:T3SS1^-^. A rise in Egr1 protein levels over time is observed only in T3SS1^+^ -infected MEFs. **C)** Immunoblot for p-Erk1/2 and total Erk1/2 in starved MEFs 45, 60, 75 and 90 minutes post-infection with T3SS1^+^ Δ*vopQ* or T3SS1^+^ Δ*vopS V. para*. T3SS1^+^ Δ*vopQ* does not induce Erk1/2 phosphorylation while T3SS1^+^ Δ*vopS* induces prolonged Erk1/2 phosphorylation. **D)** Immunoblot for total Egr1 in starved MEFs 45, 60, 75 and 90 minutes post-infection with *V. para* T3SS1^+^ Δ*vopQ* or T3SS1^+^ Δ*vopS*. No rise in Egr1 protein levels is observed in T3SS1^+^ Δ*vopQ*-infected MEFs while T3SS1^+^ Δ*vopS* causes an increase in Egr1 protein levels. Target band marked with a red line and background bands with a blue star. **E)** Immunoblot showing p-Erk1/2 and total Erk1/2 in starved MEFs infected with T3SS1^+^, T3SS1^+^ *vopQ*^S200P^, T3SS1^+^ Δ*vopQ*, and T3SS1^+^ Δ*vopQ*+p*vopQ V. para* strains for 45, 60, 75 and 90 minutes. No pulse of pERK1/2 is observed in T3SS1^+^ *vopQ*^S200P^ -infected MEFs. **F)** Immunoblot for total Egr1 in starved MEFs infected for 45, 60, 75 and 90 minutes with the same *V. para* strains as in 3E T3SS1^+^ *vopQ*^S200P^ does not trigger an increase Egr1 protein levels in infected MEFs. Target band marked with a red line and background bands with a blue star. Blots are representative of N=3 independent experiments.

We repeated the infection time course with the *V. para* T3SS1^+^ Δ*vopQ* strain and found that the T3SS1-induced pulse of ERK1/2 phosphorylation was indeed dependent on VopQ as was the increase in total Egr1 protein levels (Fig. 2C,D). When MEFs were infected with T3SS1^+^ Δ*vopS*, we observed not only an amplified induction of ERK1/2 phosphorylation and Egr1 production, but also an extended duration of ERK1/2 phosphorylation (Fig. 2C,D). This observation is consistent with previous work that found that VopS’s AMPylation of Rho GTPases inhibits host ERK1/2 and c-Jun N-terminal kinase (JNK) MAPK pathways (*15*). This early pulse of VopQ-dependent ERK1/2 MAPK signaling in MEFs is distinct from the previously reported VopQ-dependent ERK1/2 phosphorylation in Caco-2 cells during late infection time points, 3-4 hours post infection, as those assays were performed with *V. para* strains that encoded functional T3SS1, T3SS2 and TdhAS hemolysins (*19*). Taken together, these data support the model that the combined action of VopQ and VopS create a pulse of ERK1/2 MAPK signaling that is restricted to early infection timepoints resulting in the controlled expression of downstream transcription factors EGR1 and FOS.

If VopQ and VopS work together to fine-tune the host response, their co-occurrence in *Vibrio* genomes containing the T3SS1 gene cluster would be predicted to be high. To test this, we used the SyntTax server to identify all *Vibrio* strains that retained synteny in the T3SS1 gene neighborhood (Table S7). We identified 58 *Vibrio* strains representing 8 species containing the T3SS1 gene cluster and found that 91.4% of genomes containing *vopQ* also contained *vopS* (53/58) (Table S7). We found that these genes co-occur in diverse *Vibrio* species including *V. parahaemolyticus, V. diabolicus, V. antiquarius, V. campbelli* and *V. alginolyticus* (Fig. S4). Interestingly, the 5 genomes containing *vopQ* but lacking *vopS* belonged to two *Vibrio* species, *V. harveyi* and *V. tubiashii*.

### VopQ-induced pro-survival signaling is independent of endosomal deacidification

Next, we wanted to understand how VopQ could activate ERK1/2 MAPK signaling in the host. The VopQ channel de-acidifies vacuolar and lysosomal compartments, but also inhibits homotypic fusion of yeast vacuoles, a model for Rab GTPase- and SNARE-dependent fusion between the lysosome and autophagosome (*9, 20*). Mutation of serine 200 to a proline creates a mutant, VopQ^S200P^, that is still able to neutralize the vacuole or lysosome, but can no longer block fusion (*9*). This observation is likely due to reduced binding of VopQ^S200P^ to the V-ATPase (*16*). To test if VopQ’s activation of ERK1/2 MAPK signaling was caused by one or both of these functions, we exchanged the chromosomal copy of the *vopQ* gene in the *V. para* T3SS1^+^ strain with a version encoding VopQ^S200P^ creating the *V. para* T3SS1^+^ *vopQ*^S200P^ strain. We tested the cytotoxicity of this strain during MEF infection and found that the VopQ^S200P^ mutant was no less lethal than wild-type VopQ (Fig. S3A). However, unlike its parent strain, *V. para* T3SS1^+^ *vopQ*^S200P^ was unable to induce ERK1/2 phosphorylation and downstream production of Egr1 in MEFs (Fig. 2E,F). Notably, treatment of MEFs with chloroquine, a drug that prevents lysosomal acidification, is not able to induce phosphorylation of ERK1/2 (Fig. 2E). These data suggest ERK1/2 MAPK signaling is not activated by lysosomal deacidification alone and is instead dependent on VopQ’s strong physical interaction with the V-ATPase. We therefore propose two models by which VopQ could induce ERK1/2 MAPK signaling. In the first, VopQ’s inhibition of lysosome-autophagosome fusion directly activates ERK1/2 MAPK signaling. In the second, VopQ manipulates another pathway upstream of both lysosome-autophagosome fusion and ERK1/2, thereby altering these two pathways in parallel.

### VopQ-induced pro-survival ERK1/2 signaling is dependent on IRE1

To test these models, we aimed to identify a signaling pathway that could be upstream of both autophagosome-lysosome fusion and ERK1/2 MAPK signaling. The unfolded protein response has previously been linked to both of these processes through IRE1’s connections to the UPR and the ERAD pathway (*21-24*). Moreover, induction of ER stress with a proline analogue was previously shown to partially stimulate IRE1-dependent ERK1/2 activation to promote cell survival by an undefined mechanism (*25*). To examine if the UPR played a role, we tested if VopQ’s activation of ERK1/2 and downstream EGR1 production was dependent on any of the three branches of UPR: IRE1, ATF6, or PERK(*26*). We infected *IRE1*^*-/-*^ MEFs, *Atf6*^*-/-*^ MEFs, *PERK*^*-/-*^ MEFs and wild-type MEFs with *V. para* T3SS1^+^ and *V. para* T3SS1^-^ and monitored phosphorylation of ERK1/2 and accumulation of Egr1 by Western blot. While infection of *Atf6*^*-/-*^ MEFs, *PERK*^*-/-*^ MEFs, and wild-type MEFs with *V. para* T3SS1^+^ resulted in robust ERK1/2 phosphorylation 45 and 60 minutes post-infection, phosphorylated ERK1/2 was not detected in *IRE1*^*-/-*^ MEFs infected with *V. para* T3SS1^+^ (Fig. 3A,B). Consistent with this observation, accumulation of Egr1 was also not observed in *IRE1*^*-/-*^ MEFs infected with *V. para* T3SS1^+^ (Fig. 3A). VopQ was still cytotoxic to *IRE1*^*-/-*^ MEFs (Fig. S5A), consistent with the maintained cytotoxicity of VopQ^S200P^ in wild-type MEFs despite its inability to activate ERK1/2 (Fig. S3A). These data suggest that VopQ induces ERK1/2 phosphorylation by an IRE1-dependent mechanism.

**Figure 3.**
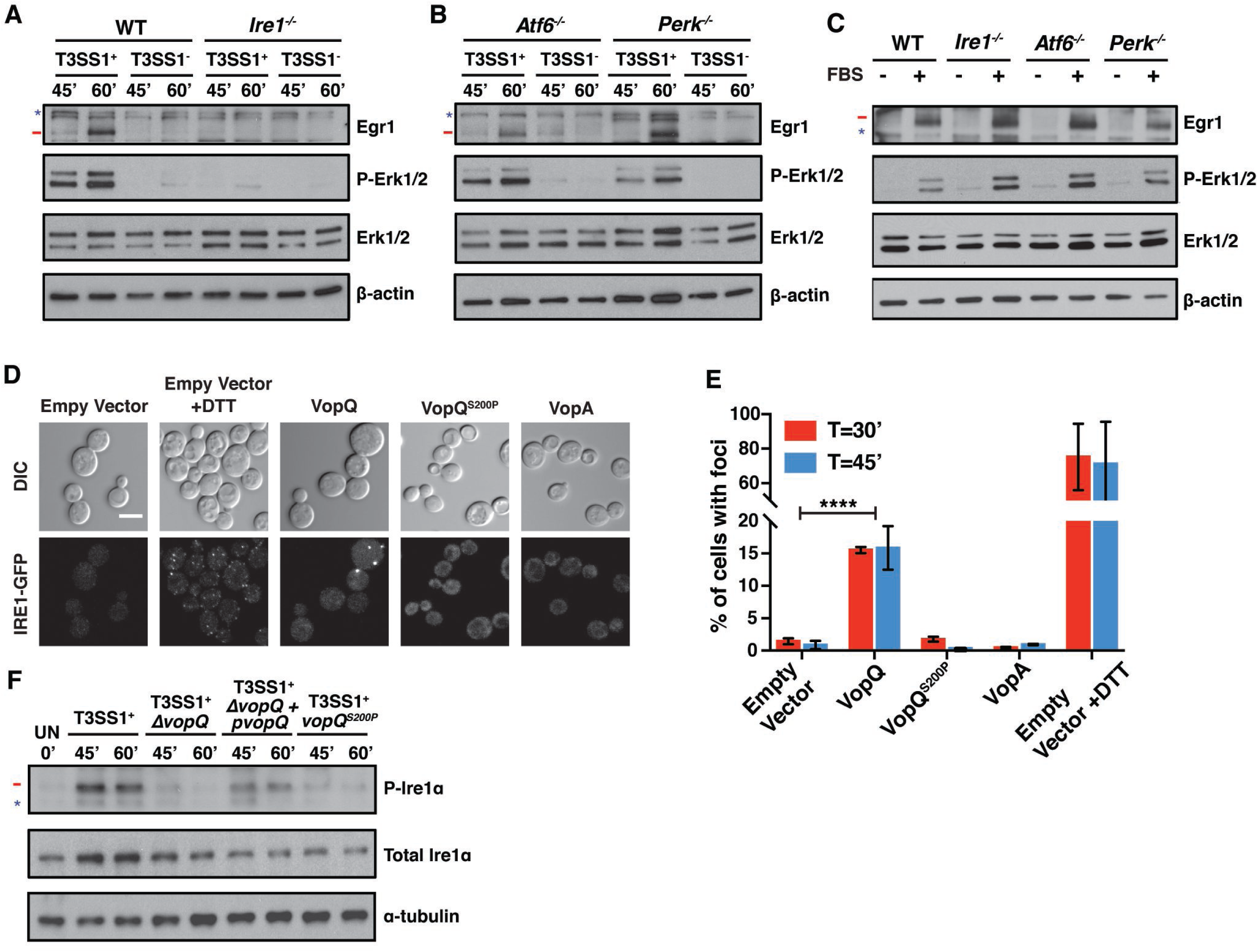
Activation of ERK1/2 MAPK signaling by VopQ is dependent on IRE1 activation. **A)** Immunoblots for total Egr1, p-Erk1/2, and total Erk1/2 in starved wild-type (WT) and *Ire1*^*-/-*^ MEFS infected for 45 and 60 minutes with *V. para* T3SS1^+^ or *V. para* T3SS1^-^. Erk1/2 phosphorylation and increased Egr1 protein levels are not observed in *Ire1*^*-/-*^ MEFS. **B)** Immunoblots for total Egr1, p-Erk1/2, and total Erk1/2 in starved *Atf6*^*-/-*^ and *Perk*^*-/-*^ MEFS infected for 45 and 60 minutes with *V. para* T3SS1^+^ or *V. para* T3SS1^-^. **C)** Immunoblots for total Egr1, p-Erk1/2, and total Erk1/2 in starved (-) or FBS-stimulated (+) WT, *Ire1*^*-/-*^, *Atf6*^*-/-*^ and *Perk*^*-/-*^ MEFS. Target band is marked with a red line and background bands with a blue star. Blots are representative of N=3 independent experiments. **D)** Representative micrograph of Ire1p-GFP cluster formation in yeast after 45 minutes of DTT treatment or effector expression. Scale bar is 5 microns **E)** Quantification of **D**. Average percent of cells (n=100) with Ire1p-GFP foci from N=3 independent experiments. Error bars represent ?SEM. *p*-values were calculated by unpaired t test (\*\*\*\**p*<0.0001).**F)** Immunoblot for p-Ire1α and total Ire1α in MEFs infected with T3SS1^+^, T3SS1^+^ Δ*vopQ*, T3SS1^+^ Δ*vopQ*+p*vopQ*, and T3SS1^+^ *vopQ*^S200P^ *V. para* strains for 45 and 60 minutes. Target band marked with a red line and background bands with a blue star. Blots are representative of N=3 independent experiments.

VopQ-mediated cytotoxicity was also conserved in the *Atf6*^*-/-*^ and *PERK*^*-/-*^ MEFs (Fig. S5B,C). Both the *Atf6*^*-/-*^ and *PERK*^*-/-*^ cell lines exhibited the same pattern of ERK1/2 phosphorylation and Egr1 accumulation upon T3SS1^+^ infection as wild-type MEFs (Fig. 3B), indicating that VopQ’s activation of ERK1/2 was specifically IRE1-dependent. Finally, we stimulated wild-type, *IRE1*^*-/-*^, *Atf6*^*-/-*^ and *PERK*^*-/-*^ MEFs with fetal bovine serum (FBS) to assess whether the well described growth factor-stimulated ERK1/2 MAPK signaling was functional in these cell lines (*27*). FBS-stimulated ERK1/2 phosphorylation and downstream Egr1 expression was observed in all cell lines (Fig. 3C). These data strongly support our model that VopQ’s IRE1-dependent activation of ERK1/2 occurs through pathway that is separate from the established growth factor-stimulated pathway mediated by Ras and Raf (*27*).

### VopQ expression results in IRE1 activation in yeast

Next we asked if VopQ activates IRE1. VopQ toxicity is dependent on the assembly c subunit of the V_o_ V-ATPase in yeast independent of the vacuolar localization of the complex. This led us led us to hypothesize that interaction of VopQ and the c subunit ring could also take place in the endoplasmic reticulum (ER) where the V_o_ complex initially forms (*16*). In addition, the ER is the source of membranes for autophagosomes, thus a block in autophagic flux may also perturb the protein to lipid ratio of the ER (*28*). We predicted that if either of these scenarios occurred, the membrane perturbations caused by VopQ may activate the UPR, which is mediated by IRE1 in yeast, either through the disruption of the ER lumen environment or through the activation of lipid bilayer stress (*29-32*). IRE1 is a type I ER-resident transmembrane protein that contains a protein kinase and an endoribonuclease domain in its cytoplasmic region (*33*). IRE1 also contains an amphipathic helix which can sense perturbations in the lipid bilayer, leading to activation and initiation of the UPR (*30*).

To test this hypothesis, we assessed the clustering of IRE1 in yeast by visualizing endogenously expressed IRE1p-GFP (20). Plasmids carrying galactose-inducible genes encoding wild type VopQ, a V-ATPase binding mutant VopQ^S200P^, and VopA were transformed into the BY4741 IRE1-GFP yeast strain. VopA is a *V. para* T3SS2 effector that kills yeast by a mechanism that is distinct from VopQ and was included as a control (*34*). Upon galactose induction serial growth assays showed that VopQ and VopA both inhibited growth in the BY4741 IRE1-GFP strain while VopQ^S200P^ and vector alone control did not (Fig. S6). We then monitored IRE1p-GFP clustering, or foci formation, at 30 and 45 minutes post galactose induction by confocal microscopy. Treatment with dithiothreitol (DTT) for the same period was used as a positive control for UPR stress. Yeast expressing VopQ, but not VopQ^S200P^ or VopA induced IRE1p-GFP foci formation (Fig. 3D). IRE1p-GFP foci formation was observed, on average, in about 70% of DTT treated cells compared to about 15% in cells expressing VopQ (Fig. 3E). This difference was expected because DTT treatment is homogeneous, whereas expression of VopQ is stochastic (*35*). Our data indicate that expression of VopQ in yeast results in IRE1 activation.

### VopQ causes induction of the IRE1 branch of the UPR of mammalian cells during infection

We next asked if VopQ also activates IRE1 in mammalian cells during infection. IRE1 is normally sequestered by BiP, but is released upon UPR activation when it then oligomerizes, *trans*-autophosphorylates and activates its endoribonuclease activity, resulting in the non-conventional splicing of XBP1 mRNA in mammalian cells (*36-38*). To determine if the IRE1 branch of the UPR was activated by VopQ in early infection time points, we measured levels of phospho-IRE1α in MEFs by Western blot at 45 and 60 minutes post infection. Our results indicate that *V. para* T3SS1^+^ induces IRE1α phosphorylation in a VopQ-dependent manner (Fig. 3F) and that T3SS1^+^ *vopQ*^S200P^ is not able to induce IRE1α phosphorylation. Taken together, these data indicate that the V-ATPase binding activity of VopQ activates the IRE1 branch of the UPR in both MEFs and yeast.

### VopQ-induced pro-survival ERK signaling is dependent on IRE1 kinase activity

Next, we wanted to further determine if the catalytic activities of IRE1 are required for VopQ-induced ERK1/2 signaling. IRE1 contains both protein kinase and an endoribonuclease domain in its cytoplasmic region (*33*). To test if the kinase or endonuclease activity of IRE1 is required for ERK signaling, MEFs were treated before *V. para* infection with KIRA6 or 4μ8c, an IRE1 specific kinase inhibitor and a potent inhibitor of IRE1 RNase activity, respectively (Fig. 4A). Strikingly, ERK1/2 phosphorylation and Egr1 protein expression were not observed in infected MEFs treated with the kinase inhibitor KIRA6. However, 4μ8c treatment appeared to have no effect on VopQ-induced ERK1/2 phosphorylation and Egr1 protein expression. Taken together, this data indicates that the pro-survival ERK signaling induced by VopQ during *V. para* infection is dependent on IRE1 kinase activity.

**Figure 4.**
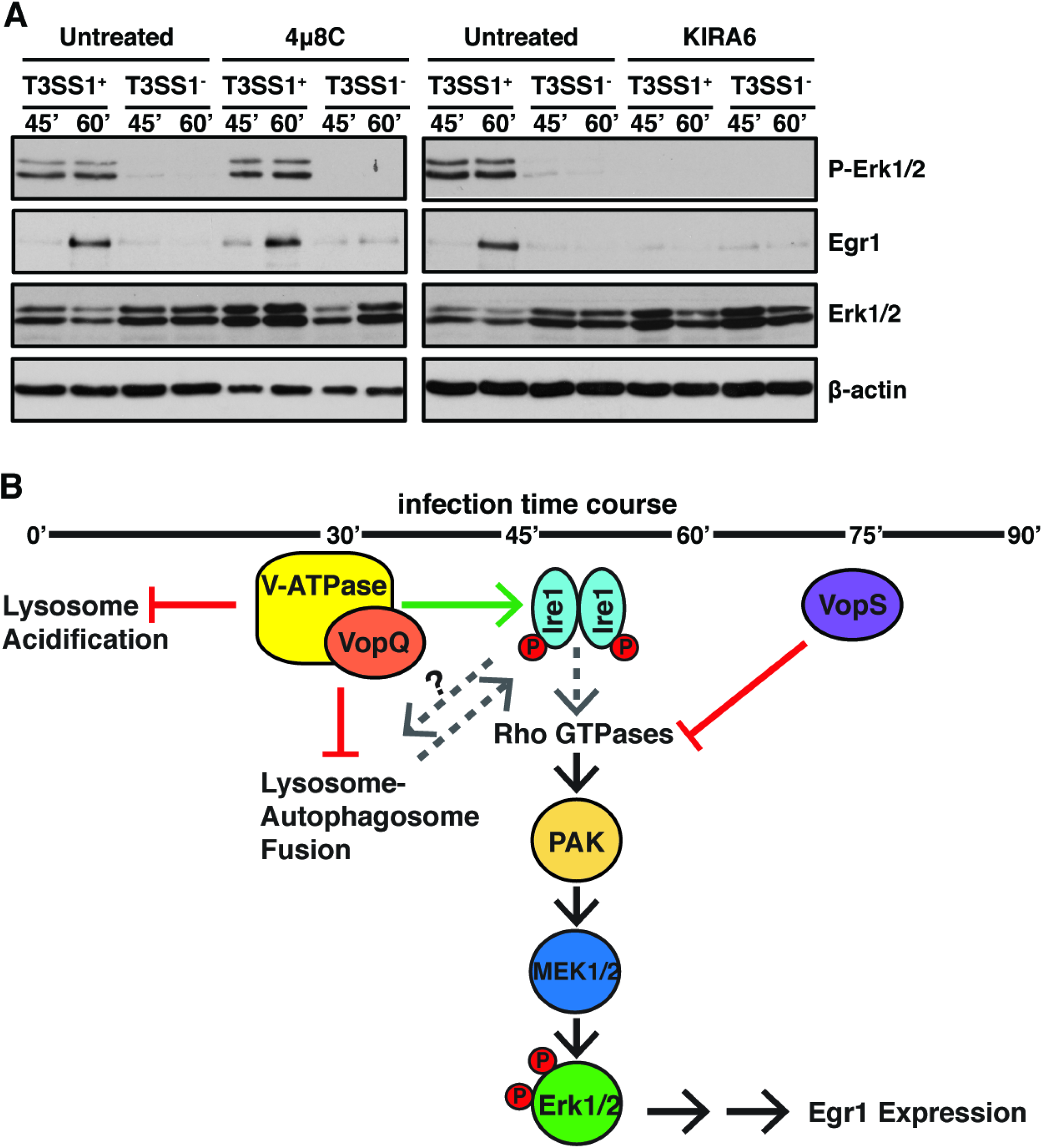
Activation of ERK1/2 MAPK signaling by VopQ is dependent on IRE1 kinase activity. **A)** Immunoblots for total Egr1, p-Erk1/2, and total Erk1/2 in starved wild-type (WT) MEFS untreated or treated with 4μ8c or KIRA6 inhibitors for 24 hours and 1 hour, respectively and infected for 45 and 60 minutes with *V. para* T3SS1^+^ or *V. para* T3SS1^-^. Erk1/2 phosphorylation and increased Egr1 protein levels are not observed in MEFS treated with KIRA6 kinase inhibitor. Blots are representative of N=3 independent experiments. **B)** Model for IRE1-dependent modulation of Erk1/2 MAPK signaling by VopQ and VopS during *V. para* infection. Green arrows indicate activation and red lines depict inhibition. Dashed lines indicate postulated connections in the model requiring future study. Specifically, it is possible that Rho GTPases may be upstream of Ire1 and how VopQ’s inhibition of autophagic flux affects Ire1 signaling and vice versa is unclear.

## Discussion

By studying the function of two effectors of the T3SS1 of the seafood-borne pathogen *V. para*, we have shown that T3SS effectors act together to systematically manipulate the host response. We observe that the effector VopQ is responsible for the T3SS1-mediated activation of cell survival and repression of cell death networks and another effector VopS is responsible for dampening this response. Of note, VopS has been established to AMPylate and thereby inactivate Rac, a known activator of MEK1/2 mediated ERK signaling(*17*). Furthermore, in *Vibrio alginolytcus*, a *Vibrio* species closely related to *V. para*, VopS was found to be required for the rapid induction of apoptosis in infected cells (*39*). The association of VopQ with the V-ATPase elicits early activation of ERK1/2 phosphorylation that is turned off by the delayed temporal action of VopS (Fig. 4B).

Manipulation of the V-ATPase to affect signaling in the cell has been previously observed with oncogenic Ras, which induces a Rac-dependent plasma membrane ruffling and micropinocytosis. However, in the case of oncogenic Ras, activation of Rac is dependent on the activity of the V-ATPase along with its relocation to the plasma membrane (*40*). Importantly, the activation ERK1/2 MAPK signaling by VopQ does not occur through the growth factor inducible Ras-mediated pathway, but instead is dependent on the kinase activity of the ER stress sensor and cell-fate executor, IRE1 (Fig. 4B). Activation of ERK1/2 MAPK signaling specifically through the IRE1-branch of the UPR by a bacterial effector has not previously been reported.

The evolutionary conservation of *vopQ* and *vopS* in *Vibrio* species retaining synteny in the T3SS1 neighborhood suggests that their concerted function is important for proper modulation of the host response. Effector pairs with opposing effects yet synergistic effects during infection are not unknown. YopJ and YopM of *Yersinia pestis* were previously described as having opposing effects on Interleukin signaling and caspase-1 processing which synergistically suppressed pro-inflammatory cytokines during infection (*41*). In *Salmonella*, the effectors SptP and SopE act as respective GTP activating proteins (GAPs) and Guanine nucleotide exchange factors (GEFs) for Cdc42 and Rac. Rapid degradation of SopE allows for the temporal regulation of Cdc42 and Rac activity during *Salmonella* infection (*42*). Furthermore, *Legionella pneumonphila* utilizes several such pairs of effectors such as SidM/DrrA and LepB, SidH and LubX, and AnkX and Lem3 (*43-49*). Together these effector pairs coordinate the establishment, maintenance and properly timed escape from the LCV environmental niche *Legionella* requires to replicate (*50, 51*).

The targeting of autophagy, UPR and MAPK signaling together is not unprecedented for pathogens. For example, the mycotoxin Patulin was found to manipulate these pathways though the inhibition of cathepsin B and cathepsin D, which leads to an accumulation of p62. Increased p62-mediated autophagy activates the PERK and IRE1 branches of the UPR through increased reactive oxygen species production. UPR activation then results in activation of ERK1/2 and phosphorylation of BAD, resulting in increased survival of host cells (*52*). Several viruses rely on the interplay of UPR and MAPK signaling for virulence as well. Dengue virus (DENV) relies on UPR activation of JNK signaling to induce autophagy and increase viral load of infected cells. Treatment with a JNK inhibitor decreased viral titers and reduced symptoms of DENV2 in mice (*53*). Recently the coronavirus infectious bronchitis virus (IBV) was found to rely on the activation of IRE1 and ERK1/2, but not XBP1 or JNK, for the induction of autophagic flux and pro-survival signaling during IBV infection (*54*).

We and others have observed that the ERK1/2 MAPK signaling can be regulated either directly or indirectly through the IRE1 branch of the UPR. However, it remains unclear if VopQ’s induction of IRE1 signaling is caused by the manipulation of autophagy or via localized perturbations in the ER membrane caused by VopQ’s interaction with assembly intermediates of the V-ATPase V_o_ subcomplex in the ER. Deciphering the epistatic relationship between VopQ’s association with the V-ATPase and activation of IRE1 will be the subject of future studies that will be important in understanding the role of the V-ATPase-UPR-MAPK feedback network in both cellular homeostasis and bacterial infection. Similarly, the interaction of VopQ with the V-ATPase at the ER should be further studied to determine if this interaction is sufficient to activate IRE1’s lipid bilayer stress response and ERK1/2 MAPK signaling independently of autophagy (*31, 55*). Other groups have hypothesized that Ire1’s interaction with the adapter protein Nck plays an important role in activation of ERK1/2 signaling upon ER stress; however, whether IRE1’s kinase or endonuclease activity are needed was unknown (*25*). Our studies indicate that the kinase activity of IRE1 is required, but the mechanism by which IRE1 kinase activity leads to ERK1/2 MAPK signaling is still poorly understood. Future experiments to dissect the role of Nck and Rho GTPase activation in this process would be a valuable addition to understanding this molecular mechanism of IRE1 induced, growth factor independent ERK1/2 MAPK signaling.

## Materials and Methods

### Bacterial strains and culture conditions

The *Vibrio parahaemolyticus* POR3 (POR1Δ*vcrD2*) and POR4 (POR1Δ*vcrD1*/*vcrD2*) strains were generously provided by Drs. Tetsuya Iida and Takeshi Honda of Osaka University. *Vibrio* strains were cultured at 30°C in MLB (Luria-Bertani broth +3% NaCl).All *V. para* strains except *V. para* T3SS1^+^ *vopQ*^*S200P*^ were from previous studies (Table S1). The T3SS1^+^ *vopQ*^*S200P*^ strain was created by cloning the *vopQ*^*S200P*^ allele (*9*) flanked by the nucleotide sequences 1 kb upstream and 1 kb downstream of vopQ (vp1680) into pDM4, a Cm^R^ OriR6K suicide plasmid. *E. coli* S17 (λ pir) was used to conjugate the resulting plasmid into the POR3 strain and transconjugants were selected on media containing 25mg/ml chloramphenicol. Bacteria were then counter-selected on 15% sucrose and insertion of the *vopQ*^*S200P*^ allele was confirmed by PCR.

### Mammalian Cell Culture

Primary Adult Dermal Fibroblasts ATCC^®^ PCS-201-01 (PHDFs) were purchased from ATCC and revived and maintained at 5% CO_2_ and 37°C in Low Serum Primary Fibroblast Media (ATCC) according to ATCC instructions. Wild-type Mouse Embryonic Fibroblasts (MEFs) were gift of Dr. Jenna Jewell and *IRE1*^*-/*-^, *PERK*^*-/-*^ and *Atf6*^*-/-*^ MEFs were kindly provided by Dr. Fumiko Urano. MEFs were maintained at 5% CO_2_ and 37°C in high-glucose Dulbecco’smodified Eagle’s medium (DMEM, Gibco) supplemented with 10% (v/v) fetal bovine serum (Sigma-Aldrich), 1% (v/v) penicillin-streptomycin-glutamine, and 1% (v/v) sodium pyruvate.

### Yeast strains and plasmids

All yeast genetic techniques were performed by standard procedures described previously (*56*). All strains were cultured in either rich (YPD: 1% yeast extract, 2% peptone, and 2% dextrose) or complete synthetic minimal (CSM) media (Sigma) lacking appropriate amino acids with 2% dextrose, 2% raffinose, or 2% galactose. Yeast were serially diluted and spotted onto agar plates to assay fitness and temperature sensitivity per standard technique. Yeast strains used in this study were BY4741 (MATa *his3Δ0 leu2Δ0 met15Δ0 ura3Δ0*) and BY4741 IRE1-GFP (MATa *his3Δ0 leu2Δ0 met15Δ0 ura3Δ0 IRE1-GFP*) (Thermo Fisher) as indicated.

Plasmids pRS416-Gal1-FLAG-VopQ and pRS416-Gal1-FLAG-VopQ^S200P^ were generated by sub-cloning pRS413-Gal1-VopQ and pRS413-Gal1-VopQ^S200P^ (previously published (*9*)) into the *BamHI* and *EcoRV* sites of pRS416-Gal1. Plasmid pRS416-Gal1-VopA-FLAG was generated by sub-cloning pRS413-Gal1-VopA-FLAG (previously published (*34*)) into the *EcoRI* and *XhoI* sites of pRS416-Gal1.

### Infection of PHDFs for RNA-sequencing and quantitative RT-PCR

For RNA-sequencing, PHDFs were seeded onto 6-well plates at a density of 1×10^5^ cells/mL and grown for 18-20 hours to ∼80% confluency. Overnight *V. para* cultures were normalized to an OD_600_ = 0.2 and sub-cultured to OD_600_ = 0.6. Bacteria were pelleted, resuspended in un-supplemented DMEM and grown at 37°C for 45 minutes to pre-induce T3SS1 expression (*14*). PHDFs were washed with un-supplemented DMEM and then infected with pre-induced *V. para* strains at a multiplicity of infection (MOI) of 10. Plates were centrifuged at 1000xg for 5 minutes to synchronize infection and incubated at 37°C, 5% CO_2_. At 90 minutes post-infection, RNAprotect Cell Reagent (Qiagen) was added to stop the infection and preserve the RNA. Cells were harvested by scraping, pooled and pellets were resuspended in RLT-plus buffer (Qiagen) and stored at -80°C. The same infection protocol was followed for qPCR experiments with PHDFs and MEFs. For MEK1/2 inhibition qPCR experiments MEFs were incubated with 10μM U0126 (Cell Signaling) or 10μM DMSO (vehicle) during and for one hour prior to infection.

### RNA Isolation and RNA-sequencing

RNA isolation was performed the same for PHDFs and MEFs. Cells were lysed with 27G^1/2^ needles and then homogenized with QiAshredder columns (Qiagen). Total RNA from triplicate experiments were purified with the RNAeasy Plus Kit (Qiagen). Quality of purified total RNA samples was determined by Agilent 2100 Bioanalyzer and only samples with RIN score 9 or higher were used. RNA concentration was measured by Qubit fluorimeter prior to library prep. 4μg of total DNAse treated RNA was run through the TruSeq Stranded Total RNA LT Sample Prep Kit from Illumina as previously described(*3*). Samples were quantified by Qubit before being normalized, pooled, and then sequenced on the Illumina Hiseq 2500 with SBS v3 reagents. Each sample was sequenced at a depth of at least 25 million 50-nucleotide single-end reads.

### Infection of MEFs for Immunoblotting

For ERK1/2 and Egr1 immunoblotting experiments, MEFs were starved in un-supplemented DMEM for one hour prior to infection/treatment to remove background growth factor-stimulated MAPK signaling. MEFs were not starved for in phospho-IRE1α experiments in order to prevent background activation of the UPR by nutritional stress. Cells were then infected at an MOI = 10 with *V. para* as described above or with treated DMEM supplemented with 10% (v/v) fetal bovine serum (FBS, Sigma-Aldrich). To inhibit IRE1 nuclease and kinase activity, cells were treated with 100μM 4μ8c for 24hrs and 100μM KIRA6 for 1hr before infection, respectively.

### Immunoblotting

MEFs were infected as described above and washed with 1X ice-cold PBS and collected by scraping at each timepoint. Collected cells were pelleted (1,000xg) and washed twice in ice-cold 1X PBS and lysed in RIPA buffer (50mM Tris, pH 8.0, 150mM NaCl, 5mM EDTA, 1% Nonidet P-40, 0.5% sodium deoxycholate, 0.1% SDS) with protease and phosphatase inhibitors (Roche Applied Science) for 20 minutes on ice. Total protein concentration of lysed supernatants was determined by Bradford assay and all samples were normalized for total protein prior to gel electrophoresis and immunoblotting. Total ERK1/2 and Phospho-ERK1/2 were detected with Cell Signaling Technologies (CST) p44/42 MAPK (137F5) and P-p44/42 MAPK T202/Y204 (197G2) primary antibodies, respectively. EGR1 was detected using CST EGR1 (15F7) and β-actin was detected by Sigma-Aldrich A2228 Monoclonal Anti-β-actin. Total IRE1α and Phospho-IRE1α were detected by CST IRE1α (14C10) and Novus IRE1α (pSer724), respectively. α-tubulin was detected by Santa Cruz α-tubulin (B-7). Secondary HRP-conjugated antibodies used were donkey anti-rabbit (GE Healthcare) and goat anti-mouse (Sigma-Aldrich).

### IRE1p-GFP Clustering assay

Yeast strains were grown to mid-log phase (OD_600_ ∼0.5) in CSM media lacking uracil with 2% raffinose. Cultures were then treated with 2% galactose, 2% raffinose, or 2% raffinose + 5mM DTT for 30 or 45 minutes. Cultures were collected, resuspended in 1X PBS, and fixed with 4% paraformaldehyde. Confocal images were acquired using a Zeiss LSM800 with Zen software. Images were processed with ImageJ (National Institutes of Health) and Adobe Photoshop CS6. For quantification, the presence of IRE1-GFP foci was scored in 100 cells per experiment over three independent experiments.

### Statistical Methods

For RNA-sequencing DE analysis statistical cutoffs were as follows: false discovery rate (FDR) ≤0.01, log_2_counts-per-million (log_2_CPM) ≥0 and the absolute value of fold change (FC) ≥1.5. For IPA pathway and biological network analysis the *p* values are presented as -log(Pvalue) and the cutoff for significance was *p* < 0.05. FDRs and *p* values are reported in Tables S2-6. For quantitative RT-PCR *p* values were calculated by one-way ANOVA and Dunnett’s multiple comparisons test. For the quantification of IRE1-GFP clustering *p* values were calculated by Student’s unpaired t-test (two-tailed). Additional Materials and methods are available as supplementary materials

## Supplementary Materials

## Materials and Methods

**Fig. S1**. Rapid cytotoxicity of *V. para* T3SS1 is dependent on the effector VopQ.

**Fig. S2**. Analysis of RNA sequencing data and RT-qPCR verification.

**Fig. S3**. Conservation of T3SS1 VopQ-dependent cytotoxicity and MEK1/2 dependent EGR1 expression in MEFs.

**Fig. S4**. Co-occurrence of *vopQ* and *vopS* in *Vibrio* strains with T3SS1 synteny.

**Fig. S5**. VopQ-mediated cytotoxicity is conserved in UPR mutant cell lines.

**Fig. S6**. VopQ but not VopQ^S200P^ inhibits growth of BY471 IRE1-GFP yeast.

**Table S1**. Bacterial strains used in this work.

**Table S2**. Complete RNA sequencing Differential Expression Data.

**Table S3**. Differential Expression of T3SS1-specific genes.

**Table S4**. IPA Canonical Pathway Analysis of Differential Expression Data.

**Table S5**. IPA Network (Disease and Biofunction) Analysis of Differential Expression Data.

**Table S6**. Expression of alternate MAPK pathway target genes

**Table S7**. SyntTax VopQ and VopS cooccurence in Vibrio genomes with T3SS1 Synteny.

## Acknowledgements

We thank Dr. Fumiko Urano for generously providing MEF *IRE1*^*-/-*^, *Atf6*^*-/-*^ and *PERK*^*-/-*^ cell lines, as well as Dr. David Ron and Dr. Randal Kaufman.

## Author contributions

RNA-sequencing experiments were performed by N.J.D. Analysis of RNA sequencing data as well as IPA network and pathway analysis was performed by M.K. and N.J.D. Quantitative RT-PCR experiments were carried out by N.J.D. MEF infections and Westerns were carried out by N.J.D, A.E.L, A.K.C., and F.S.C. Yeast strains were prepared by A.K.C. and IRE1-GFP clustering assays were performed by A.K.C and N.J.D. Synteny analysis was carried out by L.N.K. The manuscript was written by N.J.D. and A.K.C. with substantial contributions from K.O. Figures were prepared by N.J.D, M.K., A.K.C and L.N.K. The research was funded by grants to K.O., C.X. and N.V.G.

## Data and materials availability

The authors declare that the data supporting the findings of this study are available within the paper and the Supplementary Information. Complete RNA-sequencing data has been deposited on the Gene Expression Omnibus server (GSE120273). All data is available from the corresponding author upon request.

